# An α-1,3-fucosyltransferase influences thermal nocifensive escape behaviors in *Drosophila* larvae

**DOI:** 10.1101/2025.06.02.657458

**Authors:** Shannon L. Ballard, Lauren Holt, Elizabeth Stant, Ashley Arthur Moore, Luisa Zavala, Troy R. Shirangi

## Abstract

When exposed to noxious thermal or mechanical stimuli, *Drosophila melanogaster* larvae will attempt to escape by performing a stereotypical nocifensive rolling behavior. Here, we report the identification of a mutation in *Drosophila* that specifically alters the thermal nocifensive escape behaviors of larvae. We provide genetic, molecular, and histological evidence that this mutation maps to the *FucTA* gene, which encodes an α−1,3-fucosyltransferase. Our results suggest that fucosylation is important for the thermal nociception rolling behavior of *Drosophila* larvae.

## Description

*Drosophila* larvae attempt to escape noxious thermal or mechanical stimuli by bending their bodies into a c-shape and rolling 360º in a corkscrew-like fashion (Boivin et al., 2023). These nocifensive escape behaviors evolved most likely as a response to attacks by parasitoid wasps (Hwang et al., 2007). In the laboratory, c-bending and rolling behaviors can be routinely elicited by gently touching the abdomen of a larva with a soldering iron set at 46ºC (Tracey et al., 2003) (**Figure 1A, Movie 1**). In a genetic screen of mutations in *Drosophila*, we identified a mutant, *S89*, whose larvae displayed an aberrant response to noxious thermal stimuli (**Movie 2**). Compared with *w*^1118^ and *S89* heterozygous controls, larvae homozygous for the *S89* mutation exhibited deficits in the ability to complete a roll (**Figure 1B**) and perform a c-bend (**Figure 1C**) when poked with a heated probe. *S89* mutants crawled with normal speeds (**Figure 1D**) and responded normally to touch (**Figure 1E**) when compared with controls, suggesting that this mutation specifically affects nocifensive behaviors.

**FIGURE 1.**
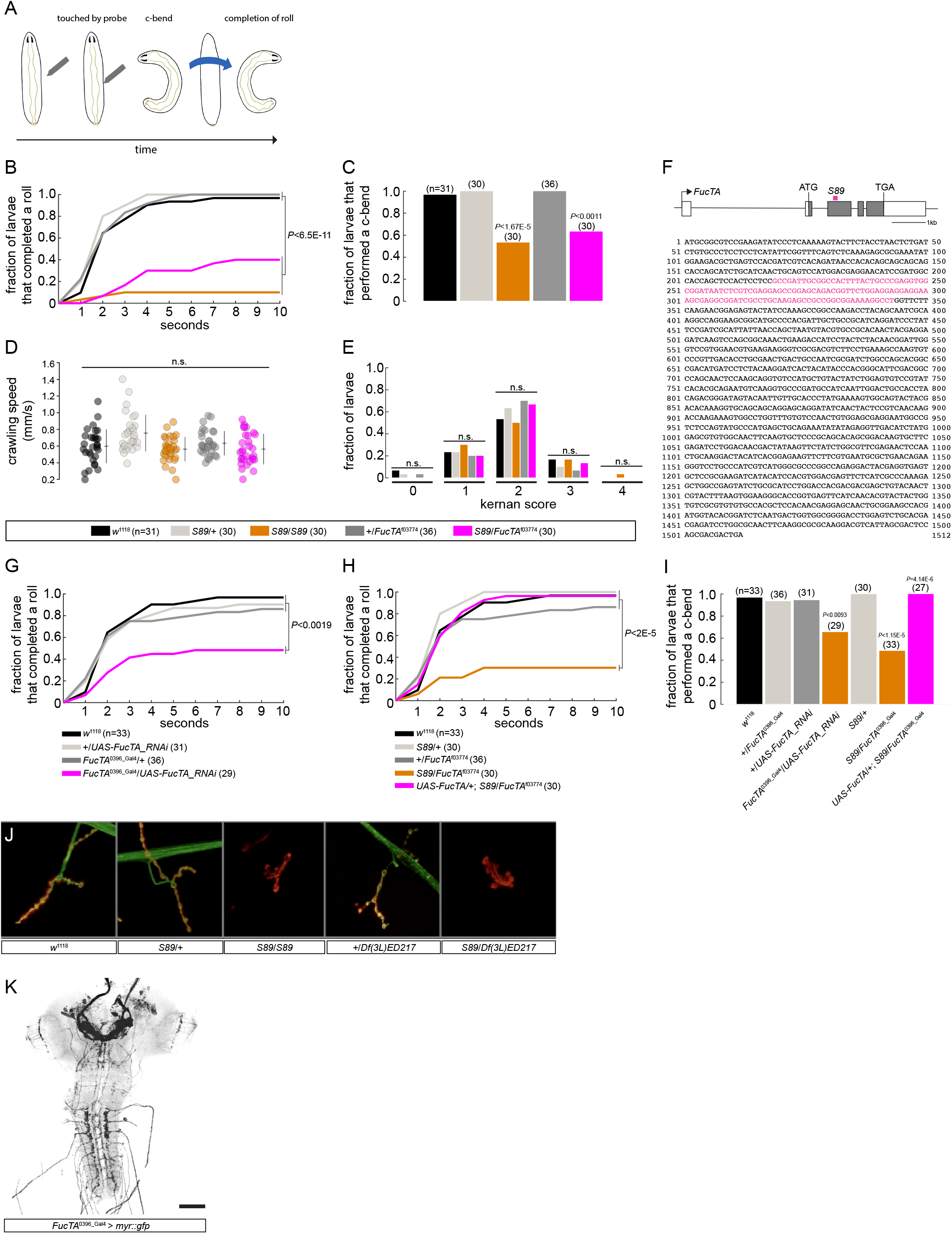
The *S89* mutation in *FucTA* causes aberrant nocifensive behaviors in *Drosophila* larvae. (**A**) Nocifensive larval behaviors such as c-bending and rolling can be elicited in the laboratory by touching the abdomen of a larva with soldering iron set at 46ºC. (**B, C**) *S89* homozygous and *S89*/*FucTA*^f03774^ mutant larvae exhibit deficits in rolling (B) and c-bending (C) behaviors after being touched on the abdomen (A2–A5) by a soldering iron heated to 46ºC. The fraction of larvae that performed a complete roll (B) or c-bend (C) during 10 seconds after being poked is shown. In (B) and (C), significance was measured using the log-rank and Fisher’s exact test, respectively. (**D, E**) *S89* homozygous and *S89*/*FucTA*^f03774^ mutant larvae exhibit normal crawling speeds (D), and responses to touch (E), as measured using the Kernan Test (Kernan et al., 1994). In (E), larvae were touched on their anterior end with an eyelash and their response was given a “kernan score” of (0) no response, (1) hesitation of the larva, (2) turning or anterior withdrawal, (3) one contractile wave backward, or (4) multiple waves backward. Significance was measured in (D) and (E) using one-way ANOVA and Fisher’s exact test, respectively. (**F**) *S89* mutants carry a 125 base-pair deletion within exon three of the *FucTA* gene. An illustration of the *FucTA* gene is shown (top). Non-coding and coding exonic sequences are colored in white and gray, respectively. The region deleted in *S89* mutants is noted in magenta. The complete cDNA sequence of *FucTA* is shown (bottom) with the 125 base-pair deletion labeled in magenta. (**G–I**) *FucTA*^0396-Gal4^-mediated expression of a UAS-regulated *FucTA* RNAi recapitulates the rolling (G) and c-bending (I) phenotypes of *FucTA* and *S89* mutants. *S89*/*FucTA*^0396-Gal4^ larvae exhibit rolling (H) and c-bending (I) phenotypes that are rescued with the presence of a UAS-regulated *FucTA* cDNA. Significance in (G) and (H), and in (I) was measured using a log-rank and Fisher exact test, respectively. (**J**) Homozygosity of *S89* abolishes anti-HRP reactivity at the fourth neuromuscular junction (NMJ) in abdominal segment 3 or 4. HRP is shown in green and the post-synaptic sites of the NMJ are shown in red using an anti-Cactus antibody. Larvae heterozygous for *S89* and a deficiency that removes the *FucTA* gene exhibit similar results. *S89* mutants display a reduction in the number of boutons at NMJ4 (data not shown) (**K**) Confocal image of a central nervous system from a *FucTA*^0396-Gal4^ > *UAS-myr::gfp* larva. GFP-expressing cells in black. Scale bar = 50 µm.

A chromosomal deficiency screen mapped the *S89* mutation responsible for the aberrant escape phenotype to an interval on the third chromosome that contains an α−1,3-fucosyltransferase encoded by the *FucTA* gene. We sequenced *FucTA*’s exons in *S89* mutants and discovered a 125 base-pair deletion within its third exon, which would substantially truncate the FucTA protein if translated (**Figure 1F**). We next performed a series of experiments that collectively provide evidence that the nocifensive phenotypes of *S89* mutants are indeed due to loss of *FucTA* activity. First, we found that the *S89* mutation is allelic to a previously characterized loss-of-function allele of *FucTA, FucTA*^f03774^ (Yamamoto-Hino et al., 2010). *S89*/*FucTA*^f03774^ larvae displayed a reduction in rolling and c-bending behaviors that was comparable to *S89* mutants (**Figure 1B, C**) and no alteration in crawling speed or touch responses compared with controls (**Figure 1D, E**). Second, RNAi-mediated reduction of *FucTA* transcripts using an enhancer-trap Gal4 allele of *FucTA, FucTA*^0396-Gal4^ (Gohl et al., 2011), with a validated UAS-*FucTA_RNAi* transgene, recapitulated the rolling and c-bending phenotypes of *S89* mutant larvae (**Figure 1G, I**), indicating that *FucTA* activity is required for nocifensive escape behavior in larvae. Third, *FucTA*^0396-Gal4^-driven expression of a UAS-regulated cDNA encoding a wild-type *FucTA* transcript rescued the nocifensive phenotypes of *S89*/*FucTA*^0396-Gal4^ mutant larvae (**Figure 1H, I**). Fourth, previous studies have shown that the neural antigen in *Drosophila* detected by antibodies against horseradish peroxidase (HRP) is a glycoprotein (Jan & Jan, 1982; Kurosaka et al., 1991; Snow et al., 1987) whose detection by anti-HRP requires *FucTA* gene function (Fabini et al., 2001; Yamamoto-Hino et al., 2010). Indeed, like *FucTA* loss-of-function mutants (Fabini et al., 2001; Yamamoto-Hino et al., 2010), larvae homozygous for *S89* lacked HRP immunoreactivity at an abdominal neuromuscular junction compared with control larvae (**Figure 1J**).

Our results indicate the *S89* mutation is a loss-of-function allele of the *FucTA* gene, and that *FucTA* activity and the process of fucosylation contribute to nocifensive escape behaviors in *Drosophila* larvae. The glycoprotein substrate(s) targeted by *FucTA* in its regulation of nociception, and where *FucTA* activity is required for larval escape behaviors are currently unknown. Muscle and motor neurons are unlikely sites, as depletion of *FucTA* transcripts using drivers that preferentially target these cells failed to produce nocifensive phenotypes (unpublished results). *FucTA*^0396-Gal4^ labels a relatively sparse number of sensory and central neurons in the larval nervous system (**Figure 1K**), suggesting that *FucTA* may contribute to nocifensive escape behaviors by functioning in at least some of these cells.

## Methods

### Drosophila husbandry and stocks

All fly stocks used in this study were maintained at 18ºC on standard cornmeal-agar-molasses media. Crosses were performed at 25ºC. The following stocks were obtained from Bloomington Drosophila Stock Center at Indiana University: *w*^1118^, *FucTA*^f03774^ (BL# 18690), *UAS-FucTA_RNAi* (BL# 51715), *FucTA*^0396-Gal4^ (BL# 64718), *Df(3L)ED217* (BL# 8074). The *S89* mutation was induced by mutagenesis with ethylmethane sulfonate and was part of a collection of ts-paralytic mutants in B. Ganeqky’s laboratory at University of Wisconsin-Madison. UAS-FucTA comprises the full-length *FucTA* sequence cloned into pUAST-attB. cDNA constructs were generated by amplification of the *FucTA* coding sequence from a full insert cDNA clone (Berkeley Drosophila Genome Project FI20174). UAS-FucTA was inserted into attP40 in a *w*^1118^ background.

### Larval nociception behavioral experiments and analyses

The larval nociception experiments were conducted similarly to the experiments performed by (Tracey et al., 2003). Briefly, third instar larvae were taken from fly food before the wandering stage. The food and larvae were placed in a petri dish with water to soften the food. Larvae were then transferred into a drop of water onto a clear sylgard-coated petri dish and given ~2 minutes to acclimate to the petri dish. The behavior of each larva was recorded (via plugable, and a standard USB microscope eyepiece camera) software before, during, and after being touched on abdominal segments A3–A6 on the left or right side with a soldering iron set to 46ºC. Once the larva initiated a nocifensive response, the soldering iron was removed from its cuticle. Otherwise, the soldering iron was held to the abdominal cuticle for 10 seconds. If there was no response after 10 seconds, the soldering iron was removed. Larval behavior was analyzed using Shotcut. The time to complete a roll was calculated by subtracting the time of initiation (the time the larva started the rolling behavior) from the time it took to complete the roll. The trachea on the dorsal side of larvae were observed to ensure that the roll was a complete 360º.

### Larval speed and touch response experiments and analyses

Third-instar crawling larvae were examined similarly to experiments performed by others (Kernan et al., 1994). Briefly, larvae were placed on a clear sylgard coated petri dish and while moving forward were touched with an eyelash tip on one side of their thoracic segments. Their behavioral response was recorded (as described above) and a score was assigned based on Figure 2 of Kernan et al. 1994. To measure larval speed, graph paper was placed underneath a clear sylgard-coated petri dish containing third-instar wall-crawling larvae. The length of one square on the graph paper was determined. A larva was recorded while moving along the dish and the distance and time was noted when the larva was moving in a straight line. The time it took the larva’s anterior-most segment to start at one line and reach the beginning of another line was noted in addition to the length it traveled based on the number of squares. The speed of the larva was then calculated.

### Immunohistochemistry

Larval CNS and NMJ dissections and stainings were conducted as previously described (Ballard et al., 2014).

## Acknowledgements

We thank Dr B. Ganeqky (University of Wisconsin-Madison) for his tutelage of S.L.B. during the initiation of this work; and Dr D. Tracey (University of Indiana-Bloomington) for helpful advice in setting up the behavioral experiments. This work was supported by internal funds from Swarthmore University to S.L.B., and a National Science Foundation grant to T.R.S (IOS-1845673).

